# Impact of Maternal Antibodies and Weaning Stress on the Replication and Transmission of Human H3N2 Influenza A in Piglets

**DOI:** 10.1101/2025.11.18.689042

**Authors:** Giovana Ciacci Zanella, Matias I. Cardenas, Carl R. Hutter, Celeste A. Snyder, Meghan Wymore Brand, Bailey Arruda, Daniel R. Perez, Tavis K. Anderson, Daniela S. Rajão, Amy L. Baker

## Abstract

Modern swine production facilitates indoor respiratory contact between human employees and pigs in their care, creating conditions for interspecies transmission of influenza A virus (IAV). Sow vaccination is routinely practiced in the U.S. to transfer maternal derived antibodies (MDA) to piglets. Weaning is a highly stressful period for piglets that requires increased human interaction. This study investigates the effect of maternal antibodies on the susceptibility of weaned piglets to a human-origin H3N2 IAV. Weaned piglets often possess mixed immunity from MDA, which may be antigenically matched or mismatched to circulating viruses. Given the repeated spillover of human seasonal H3N2 into swine, we specifically examined how matched and mismatched MDA, acquired from vaccinated sows, influenced piglet susceptibility. Additionally, we assessed the impact of weaning-related stress on the outcome of viral challenge. The H3N2 virus was generated by reverse genetics to mimic the 2010.1 H3N2 introduction from humans to swine. Challenged seeder piglets were divided by immune and weaning status. Two days post inoculation, naïve direct contact pigs were placed with seeders. IAV qRT-PCR and virus titration were performed on nasal swabs and bronchoalveolar lavage fluid to evaluate shedding and transmission kinetics. Matched MDA were effective in reducing shedding in challenged pigs and minimizing transmission to contacts. There was an increase in shedding and transmission in weaned pigs compared to littermates that remained on the sow. These results identify critical control points in production where changing practices could mitigate human-to-swine and swine-to-swine transmission to prevent establishment of novel lineages in pig populations.

**Importance:** Defining the factors that increase the susceptibility of pigs to infection with human influenza A viruses (IAV) is critical to understand why those viruses transmit to the new host. IAV is frequently detected in nursing pigs, where it was shown that maternal derived antibodies (MDA) may reduce clinical signs but may not prevent infection and transmission. Infected weaned piglets can then move viruses from the sow farm to offsite nurseries, where they can cause outbreaks with clinical disease as MDA wanes. Determining management practices that can be modified to reduce interspecies transmission of viruses to pigs is economically beneficial to the swine industry and could help define measures to prevent new spillover events. Reducing spillover of human IAV into pig populations also benefits public health by reducing genomic and phenotypic diversity in swine and the subsequent potential for zoonotic transmission.

## Introduction

Influenza A virus (IAV) is one of the most frequently detected swine respiratory pathogens and can cause outbreaks in swine herds throughout the year. Infected pigs develop acute respiratory disease, and the high morbidity in herds leads to significant economic losses for swine producers [1, 2]. Although only H1N1, H3N2, and H1N2 subtypes are endemic in pigs, numerous genetically and antigenically diverse IAV lineages and clades circulate in swine worldwide [1, 3]. This diversity arises from the virus’s ability to evolve through reassortment between different lineages and through mutations in its surface proteins genes, hemagglutinin (HA) and neuraminidase (NA).

Transmission between swine and humans also contributes to the genetic and antigenic diversity of swine IAV [3–5]. Frequent introductions of antigenically distinct human-origin strains into swine herds poses a significant barrier to infection control and the development and employment of broadly protective vaccines. Further, mixing different levels and specificity of population immunity promotes the maintenance of endemic IAV strains in U.S. commercial swine herds [3]. Influenza viruses notoriously cross species barriers, but to establish in a new host, there must be an initial inter- and intraspecies transmission period, followed by adaptation for efficient replication and transmission. Strains that are endemic in pigs can sometimes spill over to humans, leading to zoonotic infections. Swine-origin IAVs detected in humans, termed “variant viruses”, are reported often but rarely result in sustained human-to-human transmission [6, 7]. In contrast, reverse-zoonotic transmission events from humans to pigs are frequent and lead to sustained transmission and establishment of new lineages [5, 8]. All IAV viruses currently endemic in U.S. swine contain an HA gene that evolved from a human-origin virus [9–11]. When human IAV spillover into swine and continue to circulate, the internal genes of the human-origin viruses are often replaced via reassortment with genes from endemic swine IAV strains, potentially contributing to adaptation. However, the human-origin HA and/or NA segments are maintained [8]. This diversity creates a reservoir of viruses in swine that represent a zoonotic risk as they evolve and become antigenically distinct from the human seasonal strain that was introduced into swine.

One example of this phenomenon was the 2010.1 H3N2 swine IAV lineage. In 2012, genomic surveillance detected a novel reassorted IAV strain that contained human-seasonal H3 and N2 [12]. These viruses reassorted with endemic swine IAV while maintaining the human-origin H3 segment, and this lead to a new endemic lineage of swine IAV (2010.1 H3 lineage) [13]. This lineage has since become one of the most frequently detected H3 subtype lineages in U.S. swine [3]. The internal genes of the 2010.1 H3 HA either originated from a triple-reassortant H3N2 virus detected in swine populations in the 1990s (T) [14] or from the H1N1pdm09 (P) lineage [15]. Despite recurrent human-to-swine IAV introductions [5, 8, 16–18], not all have persisted and established new endemic swine lineages [5, 8, 19, 20]. The factors that facilitate the establishment of a human-origin virus in pigs are unclear but given the frequency of human H3 detection in U.S. swine [21], understanding these factors will help determine whether a single detection represents a threat of permanently altering swine IAV diversity.

The immune response to endemic and novel viruses can affect the evolutionary trajectory of a pathogen [22–24]. However, quantifying how variation in immune responses affects the frequency of interspecies transmission and subsequent viral adaptation to a new host species has not been systematically investigated. In U.S. commercial swine farms, herd vaccination is a common strategy to control IAV, with most vaccines administered to sows before farrowing so they can transfer maternally derived antibodies (MDA) to their piglets. These vaccines do not usually provide neutralizing protection against the all IAV circulating in pigs, and the heterogenous immune landscape can result in variable protection due to MDA mismatch to IAV that is circulating in different production locations [25, 26].

Stress is another critical factor to be considered in swine production that may influence virus transmission. Stress affects immune function and susceptibility to infectious agents in many species, including humans and pigs [27]. In modern swine production, weaning is one of the most stressful events in a pig’s life and is associated with immune system dysfunction by alterations in altered immune cell recruitment and proliferation [28, 29]. Weaning separates piglets from the sow, typically at three weeks of age, marking a significant change in their diet, environment, and immune development. In addition to the various stressors associated with weaning, this period is marked by increased human interaction, transportation and commingling of a diverse population of piglets from other litters with differing influenza immunity and infection statuses [30]. Stressed-weaned piglets with dysregulated immune responses may be more susceptible to viral infections [31], and consequently stressed pigs may represent a more favorable host conditions for sustained replication and transmission of non-adapted human-origin viruses.

In this study, we hypothesized that mismatched or lack of vaccination in the sows, and the stress of weaning negatively impact the piglet’s immune responses and this would facilitate replication and transmission of a human-origin IAV. Moreover, we predict that these standard management practices then facilitate the establishment of these viruses in the new host by allowing higher levels of viral replication and shedding compared to environments where optimal vaccination is in place and stress is minimized. To test this hypothesis, we generated a virus that mimicked the 2010.1 H3N2 human-to-swine spillover through reverse genetics for a pathogenesis and transmission study. This virus contained the HA and NA gene segments from a human seasonal H3N2 and the internal gene segments from endemic U.S. swine IAV representatives. This study determined how the host’s immune profile, specifically the transfer of maternal antibodies and stress, impacts the replication and transmission of IAV in novel hosts. Our work quantified the effect and interaction between matched, mismatched, and an absence of antibodies acquired from vaccinated sows and weaning stress on the susceptibility of piglets to a human-origin H3N2 IAV.

## Results

### Antibody responses and serum cortisol levels

We enrolled eleven bred sows in a vaccination program. Four sows received three doses of a homologous in-house whole inactivated virus (WIV) vaccine containing the challenge virus (hu-like H3N2rg containing the A138S mutation in the HA, H3 numbering) while three sows received a commercial heterologous IAV vaccine. The remaining four sows were not vaccinated against IAV (Figure 1). Sows farrowed and piglets were monitored immediately after birth to ensure colostrum intake. Hemagglutination inhibition (HI) assays were performed against hu-like H3N2rg to assess vaccine-induced neutralizing anti-influenza antibodies and the transfer of maternal antibodies to piglets. Following the second vaccination, sows that received the homologous whole-inactivated virus (WIV) vaccine developed significantly higher levels of antibodies than sows that received the commercial heterologous vaccine. This difference was measured in log-2 reciprocal titers and persisted until the sows were exposed to piglets that had been inoculated with the virus (Figure 2A). Naïve sows from the non-vaccinated group seroconverted after being exposed to their piglets that were challenged.

**Figure 1.**
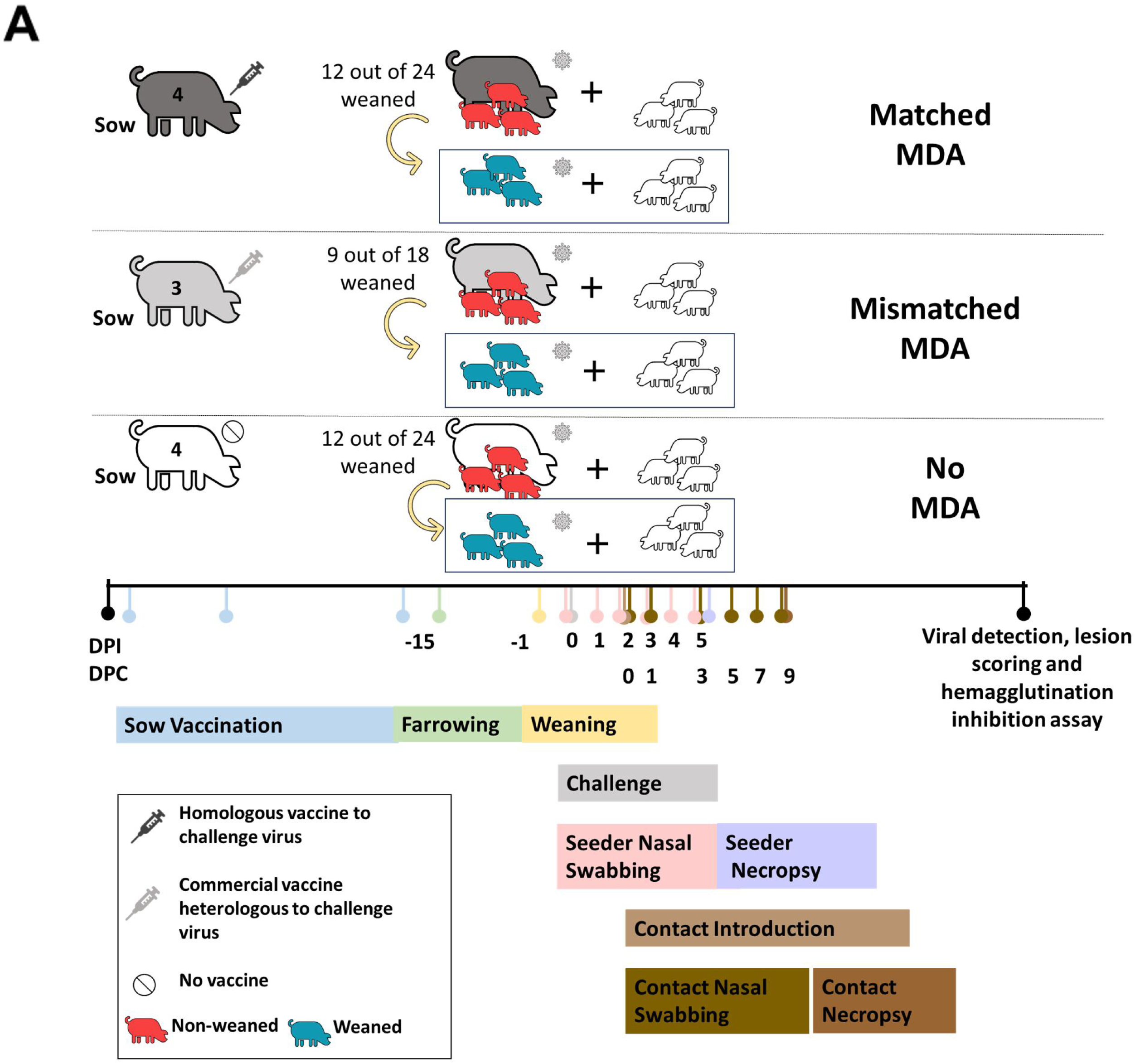
Animal study design. (A) Four sows were vaccinated with the hu-like H3N2rg whole inactivated virus vaccine, three were vaccinated with the commercial Flusure XP® and four were not vaccinated for influenza A virus. Each vaccinated sow received three intramuscular doses; the first two were within 2-week intervals before artificial insemination, and the third and last dose was administered one month before farrowing. At 15 days post-birth, piglets with matched maternal antibodies, mismatched maternal antibodies, or no maternal antibodies were further sub-divided into weaned or not weaned groups. Nasal swabs were collected daily. At 2 days post inoculation (DPI) naïve direct contact pigs were placed with seeders to evaluate transmission. Primary seeder piglets were humanely euthanized at 5 DPI and contact pigs at 9 days post-contact (DPC).

**Figure 2.**
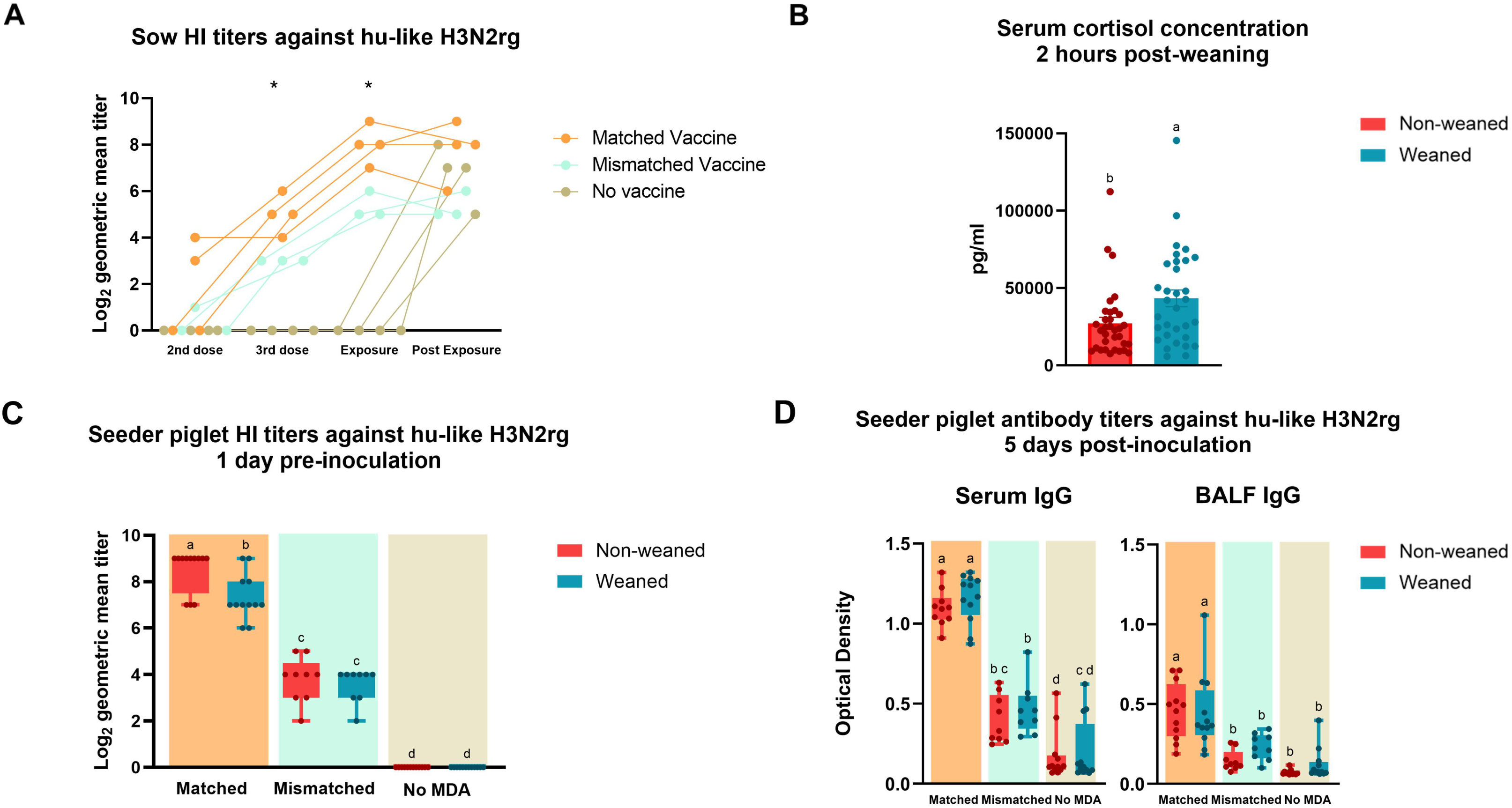
Antibody responses in sows and seeder piglets against hu-like H3N2rg. (A) Sow HI log_2_ transformed titers at the second and third vaccine doses, after exposure and post exposure to hu-like H3N2rg virus. Sows received either a matched vaccine to the challenge virus (orange), a mismatched vaccine (green), or no vaccine (brown). Data represent the individual log-2 reciprocal titers/10. (B) Serum cortisol concentrations from piglets 2 hours post-weaning assessed using a colorimetric competitive ELISA kit. Data are presented in picograms per milliliter. Piglets were either non-weaned (red) or weaned (blue). (C) Seeder piglet HI log_2_ transformed titers one day prior to inoculation. Data is stratified by maternal care group and presence or absence of maternally derived antibodies (MDA). Piglets were either non-weaned (red) or weaned (blue). (D) Seeder piglet serum and BALF IgG levels measured by ELISA five days post-inoculation. Data are presented as optical density (OD) values. Bars are showing all points. The symbol * and different lower-case letters (a,b,c) indicate statistically significant difference (p ≤ 0.05) by ordinary one-way ANOVA with Tukey’s multiple comparisons test (GraphPad Prism, GraphPad Software, La Jolla, CA).

At two weeks of age, piglets were tagged and bled for evaluation of transfer of maternally derived antibodies. Piglets were divided into groups based on maternal derived immunity and weaning status and entered the challenge and transmission phase of the study at approximately three weeks of age (Figure 1). All piglets were bled to evaluate serum cortisol levels by a commercial ELISA assay two hours post-weaning and transport of three pigs per litter. Piglets that were weaned and transported to a different containment facility showed a significant (p= 0.03) increase in serum cortisol levels when compared to piglets that were maintained with their sow (Figure 2B). Higher levels of serum cortisol in the weaned piglets demonstrated that weaning led to a physiological reaction of acute stress response. HI assays performed on piglet serum samples at weaning demonstrated they acquired MDA against hu-like H3N2rg through the ingestion of colostrum and milk (Figure 2C). Piglets born to the matched WIV vaccinated sows had robust HI titers against hu-like H3N2rg with an average of 7.9 log_2_. These piglets had significantly higher titers than those from sows vaccinated with the commercial mismatched vaccine (p= <0.0001) at weaning. As confirmed by NP ELISA (Supplemental data file) and HI assays, piglets that ingested colostrum from non-vaccinated sows did not present anti-IAV antibodies at weaning.

One-day post-weaning, seeder piglets of different immune and weaning statuses, were intranasally challenged with hu-like H3N2rg virus inoculum. 5 days post-inoculation (DPI), serum and bronchoalveolar lavage fluid (BALF) samples were collected from seeder piglets during necropsy to evaluate systemic and mucosal anti-IAV IgG (Figure 2D) and IgA (Supplemental Figure 1A) antibody levels. IgG antibodies against the hu-like H3N2rg challenge strain were evaluated by an in-house whole virus ELISA. This assay showed a similar pattern to HI pre- and post-challenge (Supplemental Figure 1B) with the matched piglet’s serum and BALF IgG antibody levels being statistically higher than both mismatched and no MDA piglet groups. IgA levels were low in serum and BALF and not statistically different between the groups. These results suggest that most IAV antibodies transferred from these vaccinated sows to their piglets and escaped the gut were the IgG isotype. Weaning did not impact systemic or mucosal anti-IAV IgG and IgA antibody levels at 5 days post-inoculation.

### Viral shedding in seeder piglets

Nasal swabs were collected daily and BALF collected at 5 DPI in the seeder piglets for the detection of IAV. No IAV was detected in the nasal swabs or BALF samples from the negative control piglets at any timepoint throughout the study (Supplemental data file). IAV was detected by qRT-PCR in nasal swabs of 15 out of 24 matched MDA piglets, whereas all nasal swabs were positive in the mismatched MDA and the no MDA piglets (Figure 3A). Seeder piglets with matched MDA had significantly lower viral titers in nasal swabs than the mismatched MDA and no MDA piglets on 1, 2, 3, 4 and 5 (p <0.0001) DPI (Supplemental data file), demonstrating that matched MDA acquired through the ingestion of colostrum from vaccinated sows protected the piglets against human-origin IAV while mismatched MDA did not. Nasal swabs from the mismatched MDA seeder piglets had significantly lower viral titers than naïve piglets without MDA on days 1 (p= 0.004), 2 (p= 0.007) and 5 (p= 0.024) post-inoculation. There was a trend for weaned piglets in the mismatched MDA group to shed higher amounts of virus than the piglets that stayed with the sow (non-weaned) and was statistically significant on day 4 DPI (p=0.033 for log 10 transformed TCID50/ml viral titer and 0.013 for Ct).

**Figure 3.**
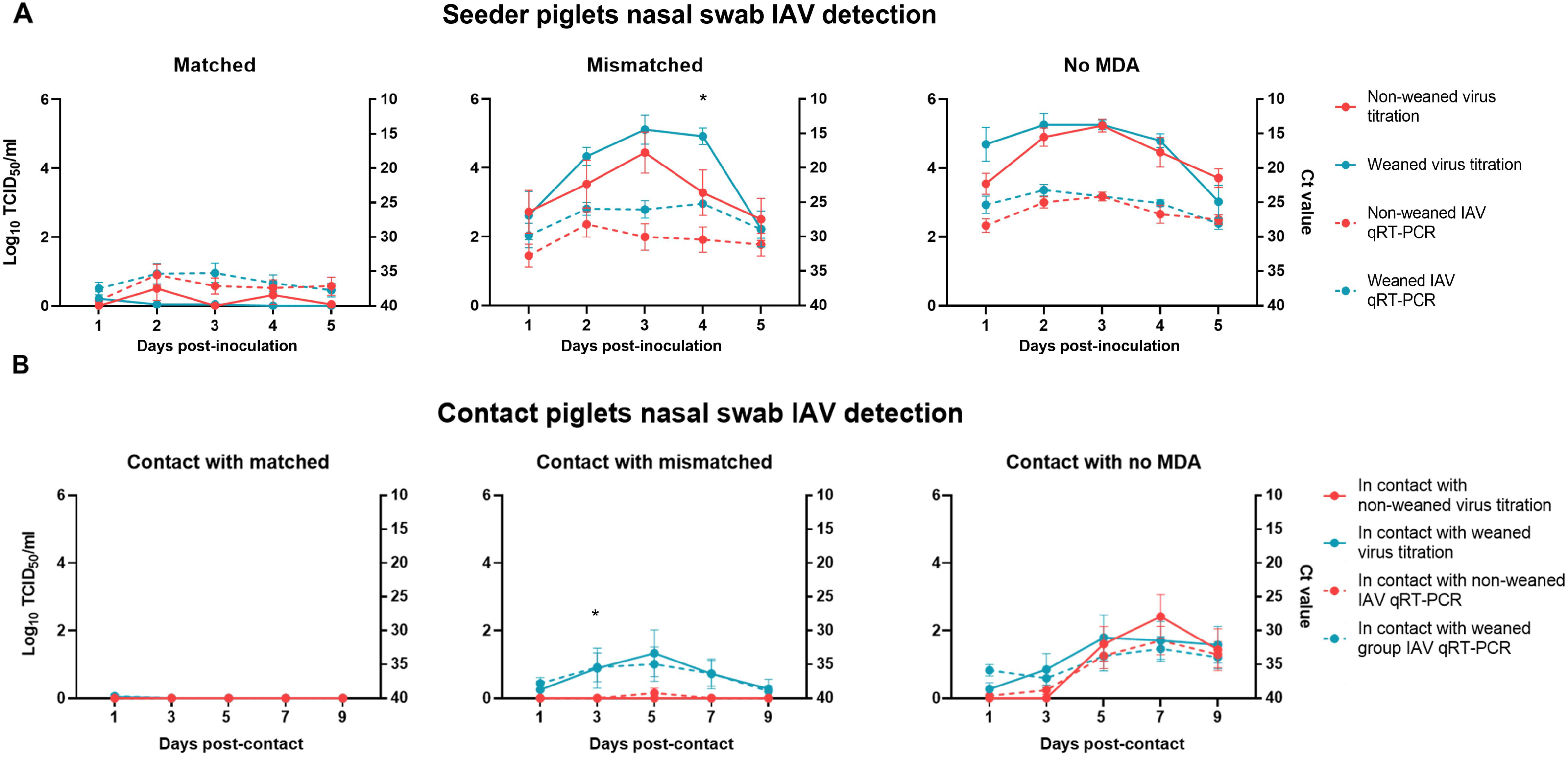
Viral detection in nasal swabs of seeder and contact piglets. (A) Influenza A virus (IAV) titers and RNA detection from nasal swabs collected from seeder piglets daily for five days post-inoculation. Virus titration results (log_10_ TCID_50_/ml, solid lines; left y-axis) and qRT-PCR Ct values (dashed lines; right y-axis) are shown for piglets non-weaned (red) and weaned (blue) across matched, mismatched and no-MDA groups. (B) IAV detection in contact piglet nasal swabs collected at days 3, 5, 7 and 9 post-contact with seeder piglets. Virus titration (log10 TCID50/ml, solid lines; left y-axis) and qRT-PCR (dashed lines; right y-axis) results are displayed for contacts of seeder piglets in the matched, mismatched and no-MDA groups. Each data point represents the mean ± standard deviation.

IAV was detected by qRT-PCR in only 1 BALF sample from piglets in the non-weaned matched MDA group but was negative for viral isolation, whereas 11 out of 18 piglets from the mismatched MDA and 18 out of 24 from the no MDA groups were positive (Supplemental Figure 1B). There was a significant difference between the amount of virus isolated from the lungs of piglets with matched MDA (group average of 0) in comparison to those with no MDA (group average of 1.2). Matched MDA were protective against IAV replication in the lower respiratory tract in addition to the upper respiratory tract. Unexpectedly, most of the seeder piglets that were positive for IAV detection in their lung samples (66.7%) were non-weaned piglets in the mismatched or no MDA groups. In addition, the non-weaned piglets from all immune statuses had significantly higher virus titers in their lungs (group average of 0.94 log) than those that were weaned (group average of 0.17) (p= 0.017). A significant relationship (p < 0.001) was found between local or systemic antibody levels (IgG and IgA) and the BALF Cts in only 35% of the samples. This indicates that antibody presence accounts for only ∼ one-third of the variation in Cts, highlighting the need to explore other factors that might be contributing to this phenomenon.

### Transmission to naïve direct contact piglets

At two days post-inoculation of the seeder piglets, naïve contact piglets were placed in nose-to-nose contact with each group of seeder piglets to assess viral transmission across different immune statuses. Only one of twenty-four MDA group contacts had virus detected by qRT-PCR in the nasal swabs on 1 DPC in the weaned group but was negative for viral isolation. (Figure 3B). Six out of eighteen mismatched MDA group contacts had IAV detected in their nasal swabs, and five of those were in contact with weaned piglets. Nineteen out of the twenty-four no MDA group contacts had IAV detected in their nasal swabs, and eleven of those were in contact with weaned piglets. There was a trend for groups in contact with weaned piglets with mismatched MDA to have more pigs shedding and at higher viral titers in nasal swabs than contacts with the piglets that remained with the sows (non-weaned) and was statistically significant on day 3 post-contact (p= 0.040). The nasal swab viral titers of the contacts placed with matched MDA were significantly lower than those of the contacts placed with no MDA piglets on days 5 (p<0.001), 7 (p<0.0001) and 9 (p<0.001) post-contact (Supplemental data file). Mismatched MDA contacts also had significantly lower nasal swab viral titers than no MDA contacts on day 7 (p<0.001) and 9 (p<0.001) post-contact.

To assess seroconversion and confirm direct transmission from seeder piglets to the contacts, serum was collected from the contact piglets at nine days post-contact and assessed by HI assay against challenge virus and NP ELISA (Table 2 and Supplemental data file). None of the contact piglets placed with the non-weaned matched piglet group had HI titers or NP antibodies detected at 9 DPC. For the piglets in contact with weaned matched MDA piglets, only one out of twelve contact piglets had a suspect HI titer of 1:20 against the hu-like H3N2rg challenge virus and this same pig had a negative NP ELISA result. The piglets in contact with mismatched MDA piglets had one out of nine non-weaned and three out of nine that were weaned with suspect titer detected by HI and only two in the weaned group had suspect values by NP ELISA. Five out of twelve piglets placed with each non-weaned and weaned no MDA groups placed with the no MDA piglets also showed signs of initial stages of seroconversion with low levels of neutralizing antibodies detected by HI. Only one piglet from the weaned group tested positive for NP antibodies and one with suspect results. There was no statistical difference in log_2_ geometric mean HI titers between the contact groups.

### Pathology

No macroscopic lesions typical of experimental IAV infection were observed in any of the lungs from seeder piglets at five DPI necropsy, consistent with a previous study [32]. In the lung sections, minimal microscopic lung lesions were present, and seeder piglet scores were comparable to the negative controls. Average microscopic lesion scores of the lungs were minimal and there was no significant difference between the scores of the MDA groups and weaning subgroups (Table 1). Microscopic tracheal lesion were more variable and average scores ranged from minimal to severe. Piglets in the matched MDA group independent of weaning status had significantly less tracheal microscopic lesions than those from the no MDA group (p=0.0001), showing that matched MDA protected piglets against tracheitis. Mismatched MDA piglets had microscopic tracheal lesion scores that were not statistically different than other groups. When accounting for weaning, there was no significant difference between the piglets overall or within the immune status group. Immunohistochemical (IHC) NP antigen staining of the trachea sections complemented the microscopic lesion scores and the IAV detection in BALF samples by demonstrating that matched and mismatched MDA significantly reduced the amount of antigen positive cells in the lower respiratory tract of piglets after challenge when compared to naïve piglets. There was trend for weaned piglets to have lower microscopic lesion and IHC scores in the trachea than non-weaned piglets inside their immunity status groups, however: these trends were not statistically significant.

**Table 1.** Macroscopic and microscopic lesions scores in the lower respiratory tract of pigs inoculated with hu-like H3N2rg.

| Group | Subgroup | Percentage of macroscopic lung lesions* | Microscopic lesion scores |  | Immunohistochemical staining score |
| --- | --- | --- | --- | --- | --- |
|  |  |  | Lung (0-20) | Trachea (0-8) | Trachea (0-4) |
| Negative controls |  | 0.00 ± 0.00<sup>a</sup> | 0.00 ± 0.00<sup>a</sup> | 0.00 ± 0.00<sup>a, b</sup> | 0.00 ± 0.00<sup>b, c</sup> |
| Matched maternal derived antibodies | Non-weaned | 0.00 ± 0.00<sup>a</sup> | 1.33 ± 0.98<sup>a</sup> | 1.08 ± 1.24<sup>a</sup> | 0.42 ± 1.16<sup>b, c</sup> |
|  | Weaned | 0.00 ± 0.00<sup>a</sup> | 1.00 ± 0.95<sup>a</sup> | 1.08 ± 1.38<sup>a</sup> | 0.00 ± 0.00<sup>c</sup> |
| Mismatched maternal derived antibodies | Non-weaned | 0.00 ± 0.00<sup>a</sup> | 0.66 ± 0.50<sup>a</sup> | 2.55 ± 1.94<sup>ab</sup> | 3.00 ± 1.73<sup>a</sup> |
|  | Weaned | 0.00 ± 0.00<sup>a</sup> | 0.77 ± 0.44<sup>a</sup> | 1.78 ± 1.99<sup>a</sup> | 1.89 ± 1.90<sup>a, b</sup> |
| No maternal derived antibodies | Non-weaned | 0.00 ± 0.00<sup>a</sup> | 1.50 ± 2.43<sup>a</sup> | 4.64 ± 2.30<sup>b</sup> | 3.6 ± 1.15<sup>a</sup> |
|  | Weaned | 0.00 ± 0.00<sup>a</sup> | 1.33 ± 0.98<sup>a</sup> | 2.75 ± 1.81<sup>a, b</sup> | 2.5 ± 1.73<sup>a</sup> |

\*Group weighted means ± standard error. Different lower-case letters within a column indicate statistically significant difference ( $p \leq 0.05$ ) by ordinary one-way ANOVA with Tukey's multiple comparisons test) (GraphPad Prism, GraphPad Software, La Jolla, CA).

**Table 2.** Distribution of positive, suspect and negative seroconversion results to influenza A virus at 9 days-post-contact across direct-contact groups.

| Group in contact with | Subgroup | Hemagglutination inhibition |  |  | NP ELISA |  |  |
| --- | --- | --- | --- | --- | --- | --- | --- |
|  |  | Positive<br>>40 | Suspect<br>20 | Negative<br>≤10 | Positive<br><0.6 | Suspect<br>0.6-0.7 | Negative<br>>0.7 |
| Matched maternal derived antibodies | Non-weaned | 0 | 0 | 12 | 0 | 0 | 12 |
|  | Weaned | 0 | 1 | 11 | 0 | 0 | 12 |
| Mismatched maternal derived antibodies | Non-weaned | 0 | 1 | 8 | 0 | 0 | 9 |
|  | Weaned | 2 | 1 | 6 | 0 | 2 | 7 |
| No maternal derived antibodies | Non-weaned | 0 | 5 | 7 | 0 | 0 | 12 |
|  | Weaned | 2 | 3 | 7 | 1 | 1 | 10 |

## Discussion

While spillover events of human IAV to pigs occur, the factors that influence the virus’s ability to infect a new host and subsequently adapt and become endemic have not been quantified. The specific human-swine interfaces where transmission events occur most frequently are also not yet identified. We hypothesized that suboptimal sow vaccination and weaning stress negatively impact the piglet’s immune responses and facilitate the establishment of human IAV in the swine host by providing a more permissive environment for virus replication and transmission. To test this hypothesis and mimic the 2010.1 H3N2 human-to-swine spillover, a virus was generated by reverse genetics and used in a pathogenesis and transmission study [32, 33]. This virus contained the HA and NA gene segments from a human seasonal H3N2 and the internal gene segments from endemic U.S. swine IAV representatives. Additionally, we introduced the A138S amino acid change in the HA that has been reported to allow swine-to-swine transmission [32]. Seeder piglets were divided by MDA immune status and weaning status and intranasally inoculated at three weeks of age. Two days post-inoculation, naïve nose-to-nose contact pigs were placed in the rooms with seeders to evaluate transmission.

Piglets without MDA were the most susceptible to human-origin IAV. They shed high amounts of virus each of all five DPI and had the highest microscopic lesion and antigen in tissue detection scores. Piglets that nursed sows that received homologous vaccination presented high levels of neutralizing MDA against the challenge virus at all blood collection time points. The matched MDA effectively reduced nasal shedding in the piglets after the challenge and significantly reduced the transmission of the human-like H3N2 to direct contacts. ELISAs demonstrated that these antibodies were primarily IgG and were detected at higher levels systemically than in BALF after challenge. Antibody levels correlated with protection against infection, replication, and lesions in the upper and lower respiratory tract. The piglets that received colostrum from sows vaccinated with the commercial heterologous vaccine had intermediate neutralizing antibody levels that were insufficient to protect from infection against the human-origin IAV. These mismatched MDA piglets shed higher amounts of virus in their nasal secretions comparable to levels in the naïve piglets. When piglets’ IAV specific MDA lack cross-reactivity to a strain that subsequently infects them, there may be a risk of vaccine-associated enhanced respiratory disease (VAERD) [34]. Though the present study included an antibody mismatch group, there was no evidence of VAERD in the macroscopic lesions scores, likely due to cross-reactivity, albeit reduced in titer.

In the U.S., 81% of large (>500 sows) breed-wean farms reported IAV sow vaccination either at pre-farrow or whole herd vaccination [35, 36]. Previous studies have shown that vaccination of gilts and sows can reduce endemic swine IAV transmission and clinical disease among the piglets. It is well established that strain-specific mass vaccination can decrease IAV shedding at the breeding herd level and the nursery [35, 37–40] and delay the time to become IAV-positive at the growing-finishing stage [41]. Our results confirm that matched MDA prevented human-to-swine transmission of IAV, while MDA derived from sub-optimally matched swine IAV vaccines did not, and the heterologous cross-protection was further reduced by weaning stress, highlighting that current vaccination programs in the U.S. may not be efficient in preventing interspecies transmission of IAV. Higher titers of IAV in the lungs of mismatched and no MDA pigs that stayed on the sow contrasted the nasal shedding results. The cause of this observation is unknown but may be due to the sows becoming infected and contributing to the viral load in the environment. It may also be due to the physiology of suckling milk from contaminated udders. Unfortunately, we did not collect nasal swabs or udder wipes to directly assess viral shedding and detection in the sows, but all sows in the non-vaccinated group, seroconverted to the challenge virus after exposure, so it can be inferred that they were infected by virus shed from the piglets and may have served as an additional viral source. Regardless, this finding also supports the use of well-matched vaccines and application strategies that maintain robust cross-reactive antibody titers in sows.

IAV circulation in pig farms enhances the risk of antigenic drift and the emergence of new genotypes [42, 43]. Piglets often act as reservoirs by maintaining and transmitting IAV within and between breed-wean farms when weaned and moved to other locations. Therefore, reducing IAV infection at weaning should decrease transmission between farms [35], lower prevalence of endemic swine IAV, and reduce risk of reassortment with newly introduced human-origin strains. We found that weaning stress increased piglet susceptibility to IAV and subsequent viral shedding and transmission in the mismatched MDA group. On day four post-inoculation, the weaned mismatched MDA piglets shed an average 42 times higher TCID_50_/ml viral titer compared with piglets in the same group that stayed with the sows. This upsurge in viral shedding could result in increased viral transmission among pigs if translated into field conditions.

In the U.S. pork industry, production practices such as weaning, transportation over long distances and mixing piglets from different litters or even source herds are embedded in modern swine agricultural practices. Our results suggest that minimal modifications to some of these production practices during the pre- and post-weaning periods can mitigate human-to-swine IAV transmission and prevent the establishment of novel lineages in U.S. swine herds. Measures like matching sow vaccines to circulating strains to increase herd immunity before farrowing, maximizing colostrum intake by all piglets in the litter to create homogenous population immunity and decreasing stressors related to weaning could decrease piglet susceptibility to human-origin IAV. Some of these production practices have also been shown to control endemic swine IAV effectively in field conditions and our data confirm that there are production practices that can also minimize interspecies transmission probability. This process, however, is challenged by a need for farm or region specific genomic surveillance data to choose representative strains for custom vaccines, and once selected the logistical and financial challenges associated a comprehensive vaccination program across multiple farms [44]. In addition, colostrum production by the sow and colostrum intake by the individual piglet are highly variable. Some of these identified risk factors may be difficult to standardize and ongoing virus circulation and evolution in the field represents a continual problem [41, 45]. Hence, maximizing sow immunity and MDA levels in piglets along with measures that reduce exposure to human-origin viruses during weaning and at the nursery should be taken as strategies for controlling human-origin IAV spillovers. Previous field research has shown evidence that workers often report to work infected with IAV of human origin that could potentially be transmitted to the pigs [46, 47]. Improving worker policies and implementing consistent and effective biosecurity practices at the swine-human interface with effective policies are the most likely, and cost-effective methods to prevent transmission. These policies could include restricting the entry of visitors, restricting access for those with influenza-like illness, enforcing farm worker sick leave, increasing IAV seasonal vaccination coverage, implementing shower-in/out policies, routine hand washing and the use of personal protective equipment (PPE) that include face masks or respirators and gloves, [40, 46, 48], and having more stringent control in farrowing rooms and for individuals that are processing piglets.

Weaning IAV-negative piglets presents an advantage to swine producers because of the benefits for pig long-term health, productivity and well-being as well as improved immune responses to IAV vaccines [48]. We provide evidence that sow vaccination and weaning stress contribute to piglet susceptibility to human-origin IAV in an experimental setting. Our results are limited to the experimental conditions, but stress is ubiquitous in agricultural swine production. In situations where piglets are stressed due to weaning, long-distance transportation, heat stress, and if any of these experiences are prolonged the impact may be more dramatic than in our controlled experiment. An additional consideration is that we artificially limited the number of pigs within experiment rooms; in a field-situation, stress may also be mediated by stocking density which tends to be much higher. Despite the limitations of our system, this study demonstrated that higher susceptibility and transmission rates due to mismatched sow vaccination and weaning stressors increased the likelihood of a human seasonal virus establishing in young pigs. Surveillance in swine must continue to be a priority for animal and public health, with priority given to specific animal–human interfaces that promote greater contact between pigs and people.

## Materials and Methods

### Virus and vaccine preparation

A reassortant H3N2 human virus (herein referred to as hu-like H3N2rg) was generated using an 8-plasmid reverse genetics system as previously described [49]. The virus contained the HA and NA gene segments from A/Victoria/361/2011 (H3N2) [VIC11], a human seasonal H3 isolate, 5 internal gene segments from a triple reassortant swine-origin H3N2 strain, A/turkey/Ohio/313053/2004 (H3N2) [OH04], and the matrix gene from the 2009 pandemic virus A/California/04/2009 (H1N1) [CA09]. Additionally, the A138S mutation in the HA (H3 numbering) was introduced by site-directed mutagenesis which has been demonstrated in previous studies to favor replication in swine lower respiratory tract cells and enhanced transmissibility in pigs. The virus was propagated in Madin-Darby canine kidney (MDCK) cells to a maximum of 3 passages following rescue, with Opti-MEM™ (Life Technologies, Waltham, MA) containing antibiotics/antimycotics and 1µg/ml of tosyl sulfonyl phenylalanyl chloromethyl ketone (TPCK)-trypsin (Worthington Biochemical Corp., Lakewood, NJ). Clarified virus from infected cell culture was diluted in phosphate-buffered saline (PBS) to a desired titer of 1 x 10^5^ 50% tissue culture infectious dose (TCID_50_)/ ml for use in the intranasal challenge.

To generate a whole inactivated virus (WIV) adjuvanted vaccine, the clarified virus was sucrose purified by ultracentrifugation at 20000 RPM for 2 hours at 4℃ (Optima™ XPN, Beckman Colter, Brea, CA) and then inactivated by UV irradiation (UV Stratalinker™1800, Stratagene; Agilent Technologies, Santa Clara, CA) for two rounds of 90 seconds duration each. Inactivation of the virus was confirmed by failure of the virus to replicate in MDCK cells. A commercial adjuvant (Emulsigen D; MVP Laboratories, Inc., Ralston, NE) was added at a 1:5 ratio. Each dose (2ml) of the WIV contained approximately 256 hemagglutination (HA) units of the virus. The heterologous vaccine was the commercially available Flusure XP® (Zoetis, Parsippany, NJ), a freeze-dried preparation rehydrated with Amphigen® that contains isolates from H1N1, H1N2 and H3N2 Clusters IV-A and B.

### Transmission study at weaning

The study design is shown in Figure 1. Eleven sows (primiparous and multiparous) were obtained from a high-health herd considered free of IAV and porcine reproductive and respiratory syndrome virus (PRRSV). Sows were housed in isolation from other animals and cared for in compliance with the Institutional Animal Care and Use Committee (IACUC) of the National Animal Disease Center (NADC). Four were vaccinated with the hu-like H3N2rg WIV, three were vaccinated with the commercial Flusure XP® and four were not vaccinated for IAV. Each vaccinated sow received three intramuscular doses; the first two were within 2-week intervals before artificial insemination, and the third and last dose was administered one month before farrowing. All sows and gilts received a dose of Farrowsure® Gold (Zoetis, Parsippany, NJ) before entering breeding protocol to prevent reproductive failure. Sows were moved to farrowing crates in animal biosafety level 2 (ABSL-2) one week before their due dates and delivered their piglets without surgical intervention.

Piglets were monitored carefully immediately after birth to ensure colostrum intake. Piglets were tagged and bled for evaluation of transfer of maternally derived antibodies at two weeks of age. All except three piglets per sow were weaned at three weeks of age and moved to a different animal building. Three piglets per sow were assigned to weaned seeder groups. Blood was collected two hours after weaning and transport to evaluate serum cortisol levels. Pigs were demonstrated to be free of influenza virus before challenge by nasal swab sampling (MagMax Viral RNA isolation kit and VetMAX™ Gold SIV Detection Kit, ThermoFisher Scientific, Waltham, MA). Those born to non-vaccinated sows and gilts were shown by an IAV nucleoprotein (NP) blocking ELISA (Swine Influenza Virus Antibody Test, IDEXX, Westbrook, ME) to be free of anti-influenza virus antibodies. All animals were housed in ABSL-2 containment during the challenge phase of the study and cared for in compliance with the IACUC of the NADC. One-day post-weaning, seventy-two three-week-old seeder piglets of different immune and weaning statuses, were intranasally challenged with a 1 x 10^5^ TCID_50_/ ml dose of the hu-like H3N2rg virus inoculum. At 2-days post-inoculation (DPI) seventy-two three-week-old naïve contacts were placed in direct nose-to-nose contact with each group of challenged piglets.

Nasal swabs were collected from seeder piglets on days 1-5, DPI, 1, 3, 5, 7, and 9 DPC in the naïve contacts. Intranasally challenged seeder piglets were bled and humanely euthanized at 5 DPI and contacts at 9 DPC with a lethal dose of pentobarbital (Fatal Plus; Vortech Pharmaceuticals, Dearborn, MI). Lungs were aseptically removed, evaluated for macroscopic lesions, and lavaged with 50 ml of MEM containing 1% bovine serum albumin (BSA) to obtain bronchoalveolar lavage fluids (BALF). A section of the right middle or affected lung lobe and distal trachea were collected and fixed in 10% buffered formalin for histopathologic examination and scoring.

### Serology and viral detection

Serum and BALF samples collected throughout the study were evaluated for anti-influenza antibodies either by the commercial NP ELISA kit (Swine Influenza Virus Antibody Test, IDEXX, Westbrook, ME) (serum only), an isotype-specific IgG and IgA ELISA (serum and BALF) and or by HI assays (serum only). For the commercial NP ELISA assay, samples were tested according to the manufacturer’s instructions. For the isotype-specific IgG and IgA ELISA, samples were tested as previously described with modifications [50]. Modifications were as follows: the serum heat inactivation time was reduced to 10 minutes and the secondary antibodies used were Goat anti-Pig IgA and IgG Heavy Chain Antibody HRP Conjugated (Bethyl labs and Fortis® Life Sciences, Boston, MA). The influenza antigen used to coat the isotype-specific IgG and IgA ELISA plates was the challenge virus hu-like H3N2rg. For the HI assays, sera were incubated with receptor destroying enzyme (RDE) overnight at 1 sera: 3 RDE in saline ratio. After RDE treatment sera was inactivated at 56℃ for 30 minutes, then incubated with 20% kaolin in PBS solution for 20 minutes, adsorbed with 0.5% and subsequently 100% turkey red blood cells (RBCs) for 20 minutes each to further remove non-specific hemagglutinin inhibitors and natural serum agglutinins. Turkey RBCs and virus dilutions were treated with neuraminidase (NA) inhibitor oseltamivir to prevent NA-mediated agglutination. Sow sera were tested against hu-like H3N2rg antigen and a surrogate Flusure XP® strain A/swine/New York/A01104005/2011 [NY/11] (Supplemental data file). At the same time, piglet sera were tested against only the challenge virus hu-likeH3N2rg. HI assays were performed using standard techniques, and titers transformed by log2 scale and geometric group means calculated [3].

Serum cortisol concentrations from piglets 2 hours post-weaning were assessed using a colorimetric competitive ELISA kit (Enzo Life Sciences, Inc., Farmingdale, NY, USA) according to the manufacturer’s instructions with the addition of the steroid displacement reagent in a 1: 99 ratio and an initial serum dilution of 1: 8. Plates were read by SpectraMax M5 Multi-Mode Microplate Reader (Molecular Devices, LLC., CA, USA). Cortisol levels were calculated using a standard curve established from cortisol kit standard samples.

All nasal swabs and BALF samples were tested for IAV RNA by RT-rtPCR (MagMax Viral RNA isolation kit and VetMAX™ Gold SIV Detection Kit, ThermoFisher Scientific, Waltham, MA). following previously validated standard protocols [51]. Samples with cycle threshold (Ct) values below 37 were subjected to virus isolation in 48-well plates containing confluent MDCK cells, in infection media and cultured at 37℃ for 72 hours. Plates were fixed when the cytopathic effect reached 80% of cells on the positive control well and stained for immunocytochemistry (ICC) staining as previously described [52]. Virus isolation-positive nasal swabs and BALF samples were tittered in 96-well plates of confluent MDCK cells in 100µl 10-fold serial triplicate dilutions. At 48 hours post-inoculation, plates were fixed and stained [52]. Titers were calculated for each sample as TCID_50_ per ml and transformed to log_10_.

### Pathology

During necropsy, the percentage of affected surface area per lung lobe was recorded and used to calculate a weighted macroscopic lung lesion score [53]. Tissues fixed in 10% buffered formalin were transferred to 70% ethanol after 48 hours. Routine histologic procedures were used to process lung and tracheal tissues, and the slides were stained with hematoxylin and eosin. Microscopic lesions were evaluated and scored by a veterinary pathologist blinded to a group with parameters previously described [54–56]. Immunohistochemical (IHC) staining was carried out manually on 5-µm thick tracheal sections, using a primary antibody that targets the IAV nucleoprotein (NP) as previously described [57]. IHC scores (0-4) were based on the amount of antigen positive cells per scored section with 0: none, 1: 1-5, 2: 6-10, 3: 11-20 and 4: >21 cells.

### Statistical Analysis

Macroscopic and microscopic pneumonia scores, HI titer log2 geometric means, log10 transformed nasal swab, and BALF viral titers were analyzed using analysis of variance (t-test (for 2 groups) and ordinary one-way ANOVA (for 3 or more groups) with Tukey’s post-hoc multiple comparisons test of parameters with statistical differences; GraphPad Prism, GraphPad Software, La Jolla, CA). Comparisons were made between the challenged groups at the same time point using a 5% significance level (p-value <0.05) to indicate statistically significant differences. Linear regression was performed between antibody quantities and CT using the lm() function in R (R Development Core Team 2025). The test was performed on the whole dataset and then on the with swine weaned groups separately.

## Acknowledgements

We thank Katharine Young, Nick Otis, Daniel Moraes, Janice Reis Ciacci Zanella and Phillip C. Gauger for their laboratory assistance and NADC Animal Resources Unit caretaker staff for support with the animals. We gratefully acknowledge pork producers, swine veterinarians, and laboratories for participating in the USDA Influenza A Virus in Swine Surveillance System. Funding was provided in part by NIH NIAID CEIRR (#75N93021C00015), National Pork Board (Project #21-085), USDA National Institute of Food and Agriculture (grant no. 2020-67015-31563/project accession no. 1022827), and USDA-ARS (ARS project number 5030-32000-231-000D). This research was supported in part by an appointment to the Agricultural Research Service (ARS) Research Participation Program administered by the Oak Ridge Institute for Science and Education (ORISE) through an interagency agreement between the U.S. Department of Energy (DOE) and the U.S. Department of Agriculture (USDA). ORISE is managed by ORAU under DOE contract number DE-SC0014664. All opinions expressed in this paper are the author’s and do not necessarily reflect the policies and views of USDA, DOE, or ORAU/ORISE. USDA is an equal opportunity provider and employer.

## Data availability statement

Clinical data associated with this study are available for download from the USDA Ag Data Commons: DOI: 10.15482/USDA.ADC/30043813.

---

**Supplemental Figure 1.** (A) Seeder piglet HI log2 transformed titers five days post-inoculation. Data is stratified by maternal care group and presence or absence of maternally derived antibodies (MDA). Piglets were either non-weaned (red) or weaned (blue). (B) Seeder piglet IgA levels in serum and BALF samples collected at five days post-inoculation, determined by ELISA. Data are shown as OD values for each group. Non-weaned group (red) and weaned group (blue). Different lower-case letters (a,b,c) indicate statistically significant difference (p ≤ 0.05) by ordinary one-way ANOVA with Tukey’s multiple comparisons test (GraphPad Prism, GraphPad Software, La Jolla, CA). (C) Seeder piglet BALF virus titration (log10 TCID50/ml; solid filled box; left y-axis) and qRT-PCR results (open box; right y-axis) at 5 DPI. Non-weaned group (red) and weaned group (blue). Numbers above the error bars show the number of positive pigs/total pig numbers in the group.

